# Graphene Quantum Dots Mitigate Oxidative Stress in Bacteria

**DOI:** 10.64898/2026.05.08.723706

**Authors:** Juhee Kim, Stephanie N. Bartholomew, Wade F. Zeno

**Affiliations:** Mork Family Department of Chemical Engineering and Materials Science, University of Southern California, Los Angeles, CA 90089, United States

**Keywords:** graphene quantum dots, microbial protection, oxidative stress, reactive oxygen species, bacterial growth, lipid oxidation, glutathione redox homeostasis

## Abstract

Manufacturing and storage processes can expose microbes to oxidative stress, reducing viability and limiting their use in biotechnological applications. Here, we evaluate graphene quantum dots (GQDs) containing hydroxyl and carboxyl groups as protective additives that mitigate peroxide-induced oxidative stress in Escherichia coli. GQDs did not adversely affect bacterial growth under basal conditions and restored growth in the presence of hydrogen peroxide. Using the membrane-partitioning fluorescent probe C11-BODIPY, we found that GQDs reduced peroxide-induced oxidation in bacterial membranes. We further used redox-sensitive roGFP2 probes to monitor intracellular oxidative stress and found that GQDs suppressed intracellular hydrogen peroxide accumulation and attenuated disruption of glutathione redox homeostasis. Together, these results show that GQDs protect bacteria by limiting peroxide-driven oxidative damage at both membrane and intracellular levels. This work supports the potential use of GQDs as protective additives for microbial formulations that are susceptible to oxidative stress.

## 1. INTRODUCTION

Microbes are utilized in various fields, such as probiotics,^1^ agriculture,^2, 3^ enzyme and supplement production,^4, 5^ and waste treatment.^6^ There is also growing interest in the applications of bacteria as therapeutic agents^7^ and in drug delivery systems.^8, 9^ During the manufacturing processes for these various applications, microbes are frequently exposed to harsh conditions that induce oxidative stress, such as drying,^10^ lyophilization,^11, 12^ and prolonged periods of storage,^13, 14^ ultimately resulting in decreased microbe viability. Oxidative stress can arise when reactive oxygen species (ROS) accumulate from environmental and intracellular sources. One particularly relevant oxidant is hydrogen peroxide (H_2_O_2_), which can be generated through photochemical reactions in aqueous solutions.^15^ Because bacterial cell membranes are permeable to H_2_O_2_,^16, 17^ extracellular H_2_O_2_ can enter cells and form hydroxyl ROS through Fenton and Fenton-like reactions, contributing to intracellular oxidative stress.^18^ ROS are also produced endogenously as byproducts of aerobic metabolism.^19, 20^ Together, these ROS can damage cells through lipid peroxidation, DNA damage, and protein damage, including reactions with sulfur-containing amino acid residues.^21, 22^

To protect microbes during manufacturing and storage, prior studies have incorporated additives such as sugars, polymers, proteins, and antioxidants into protective formulations.^23–25^ These additives are generally included to preserve microbial viability by reducing processing- or storage-associated stress. More recently, metal-phenolic networks have also been explored as protective coatings that improve microbial tolerance to oxidative environments.^26–28^ Together, these approaches highlight the value of engineering the local chemical environment around microbes to preserve viability. In this context, graphene quantum dots (GQDs) represent a potential alternative class of protective additive because of their reported water solubility, chemical stability, and biocompatible behavior in biological systems.^29–31^

GQDs are graphene-based nanomaterials with an average diameter of less than 20 nm and are composed of graphitic basal planes with oxygen-containing functional groups.^32, 33^ Due to their high biocompatibility and low toxicity, GQDs have been studied extensively in biomedical applications, and have been shown to alleviate oxidative stress.^34–36^ Based on their specific structure, GQDs can scavenge radical ROS,^37, 38^ protecting biological molecules such as lipids from oxidative stress.^39^ Although GQDs have been widely explored for their antioxidant behavior, their interactions with microbes have more commonly been studied in the context of antimicrobial activity, rather than microbial protection. For example, light-activated GQDs have been used as photothermal^40, 41^ and photodynamic^42, 43^ antimicrobial materials, where bacterial inhibition is driven by heat generation or ROS production, respectively. Other studies have enhanced antimicrobial activity by modifying GQDs with nitrogen heterocycles that damage bacterial membranes,^44^ doping GQDs with nitrogen,^45^ or conjugating GQDs to antimicrobial compounds such as curcumin.^46^ In contrast, unmodified GQDs do not necessarily exhibit intrinsic antimicrobial activity. GQDs synthesized from various precursors showed no significant antimicrobial effect or growth inhibition in the absence of light irradiation or surface modification.^47–49^ These collective works suggest that GQDs may interact with microbial systems without inherently compromising viability, raising the possibility that appropriately formulated GQDs could instead be used to protect microbes from oxidative stress.

In this study, we investigated the ability of GQDs to protect Escherichia coli (*E. coli*) from oxidative stress. GQDs at concentrations up to 50 μg/mL did not affect bacterial growth rate and even enhanced bacterial growth after prolonged incubation. Under oxidative stress conditions, GQDs restored H_2_O_2_-suppressed bacterial growth. Using the membrane-partitioning fluorescent probe C11-BODIPY, we measured the extent of lipid oxidation in bacterial cell membranes. The fraction of oxidized C11-BODIPY increased significantly following the addition of H_2_O_2_, while co-incubation with GQDs inhibited membrane oxidation. Furthermore, intracellular levels of H_2_O_2_ and glutathione redox homeostasis were monitored using redox-sensitive roGFP2 probes to investigate the effects of oxidative stress on intracellular redox balance. GQDs inhibited the accumulation of H_2_O_2_ inside the cells and attenuated the subsequent formation of glutathione disulfide, mitigating oxidative stress. Given the antioxidant ability of GQDs in living systems, our results support the potential application of GQDs as protective agents that shield bacteria from oxidative stress.

## 2. EXPERIMENTAL SECTION

### 2.1. Materials

BODIPY™ 581/591 C11 (D3861) was purchased from ThermoFisher Scientific. H_2_O_2_ (Hydrogen peroxide, 30%, HX0635) was purchased from Millipore Sigma. pQE-60 roGFP2-Orp1-His^50^ (Addgene plasmid # 64976; RRID:Addgene_64976) and pQE-60 Grx1-roGFP2-His^51^ (Addgene plasmid # 64799; RRID:Addgene_64799) were gifts from Tobias Dick.

### 2.2. GQD Synthesis and Characterization

GQDs were synthesized from carbon fiber through a thermo-oxidative cutting process.^52^ Detailed synthesis method and characterization have been described previously.^39, 53, 54^ Briefly, GQDs used in this study had an average diameter of 13.0±8.3 nm. Structural characterization confirmed the presence of graphitic basal planes and oxygen-containing functional groups such as hydroxyl and carboxyl groups.

### 2.3. Bacterial Culture

*E. coli* (BL21) was cultivated using conventional bacterial handling methods. Briefly, cells were grown on an LB agar plate overnight, and 7 mL of 2xYT media was added to the plates to scrape and suspend cells in solution. In a cuvette, 12 μL of cells were diluted in 2xYT media to a final volume of 2 mL. To measure the effect of GQDs on the growth rate, 10–50 μg/mL of GQDs were added. To induce oxidative stress, a final concentration of 4.5 mM H_2_O_2_ was added to the media. For cells that were treated with both GQDs and H_2_O_2_, GQDs were added before H_2_O_2_. Cells were incubated at 37 °C and the optical density at 600 nm wavelength (OD600) was measured hourly up to 6 hours using a spectrophotometer (Thermo Scientific NanoDrop One^C^). After overnight incubation, the OD600 was measured hourly during the final 22–24-hour period. When measuring the OD600 of cells incubated with GQDs, measurements were blanked using GQDs dissolved in 2xYT media at the corresponding concentration.

### 2.4. Microbial Membrane Oxidation

C11-BODIPY stock solutions were prepared in DMSO at a final concentration of 2 mM. *E. coli* was grown in 2xYT media at 37 °C until the OD600 reached 1.2–1.3. Cells were diluted two-fold with 2xYT media and incubated with C11-BODIPY (final concentration: 10 μM) at 37 °C for 10 minutes. Cells were then incubated with 4.5 mM H_2_O_2_, 50 μg/mL GQDs, or both. For the cells treated with both GQDs and H_2_O_2_, GQDs were added prior to H_2_O_2_. Before adding H_2_O_2_ or GQDs, the initial emission spectra of C11-BODIPY were acquired in triplicate in the range of 500–650 nm by exciting at 488 nm, using a spectrofluorometer (Jasco FP-8500). After cells were treated with H_2_O_2_ or GQDs, the emission spectra were acquired every 10 seconds for 20 minutes at 37 °C.

### 2.5. Confocal Microscopy

Imaging wells were created by placing silicone gaskets on top of Hellmanex-cleaned glass coverslips, as described previously.^55, 56^ In each imaging well, a PLL-PEG-biotin complex was added to passivate the glass surface, and the wells were rinsed with PBS to remove excess reagent. Imaging wells were then rinsed with cells that were incubated with C11-BODIPY as described in the “2.4 Microbial Membrane Oxidation” section. Using a STELLARIS 5 laser scanning confocal microscope (Leica Microsystems) and a 488 nm excitation laser, emission of C11-BODIPY was detected. The detection range was 570–620 nm and 500–540 nm for the unoxidized and oxidized states of C11-BODIPY, respectively. Imaging was conducted at room temperature.

### 2.6. roGFP2 Probe Expression and Fluorescence Measurements

*E. coli* was transformed with either pQE-60 roGFP2-Orp1-His or pQE-60 Grx1-roGFP2-His using standard bacterial transformation methods. Microbes were grown in 2xYT media with 100 μg/mL Ampicillin at 37 °C until the OD600 reached 0.5–0.6. To express proteins, 100 μM IPTG was added and incubated overnight at 20 °C. After expression, cells were washed with PBS and resuspended in PBS to a final OD600 of 0.60–0.65 and diluted two-fold to acquire excitation spectra. The initial excitation spectra were acquired in triplicate prior to adding GQDs or H_2_O_2_. The excitation intensity was measured over the range of 350–499 nm at the fixed emission wavelength of 510 nm, using a spectrofluorometer (Jasco FP-8500). Cells expressing roGFP2-Orp1 were treated with 0.1 mM H_2_O_2_, 50 μg/mL GQDs, or both. Cells were then incubated for 10 minutes at 25 °C and excitation spectra were acquired in triplicate. Cells expressing Grx1-roGFP2 were treated with 0.5 mM H_2_O_2_, 50 μg/mL GQDs, or both. After adding GQDs or H_2_O_2_, the excitation spectra were acquired every 10 seconds for 10 minutes at 25 °C.

## 3. RESULTS AND DISCUSSION

### 3.1. Impact of GQDs on bacterial growth

To determine whether GQDs influence bacterial growth under basal conditions, we monitored the growth of *E. coli* in the presence of 10, 25, and 50 μg/mL GQDs and compared these conditions to an untreated control (Figure. 1a). Over the first 6 hours, all cultures exhibited similar growth behavior, indicating that GQDs in this concentration range do not impair bacterial proliferation. After overnight growth, cultures containing GQDs reached modestly higher OD600 values, suggesting that GQDs are well tolerated and may slightly elevate the final culture density. One possible explanation for this increase is that GQDs reduce oxidative damage that accumulates during aerobic growth, allowing cells to reach a somewhat higher stationary-phase density. Regardless of the underlying mechanism, these results show that GQDs do not adversely affect *E. coli* growth at the concentrations tested here.

**Figure 1.**
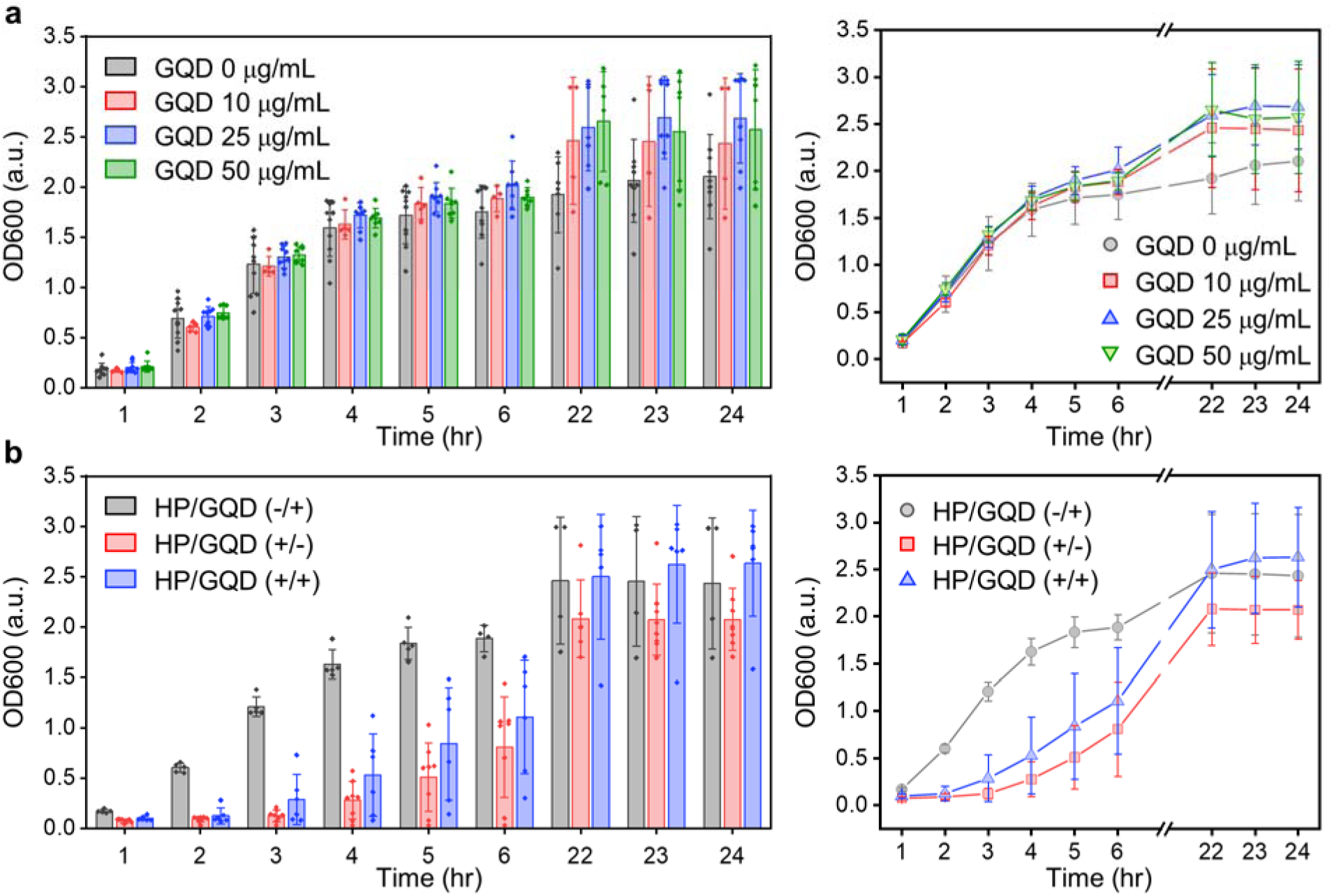
GQDs mitigate oxidative-stress induced growth inhibition in *E. coli*. (**a**) The measured OD600 of bacteria over time in the presence or absence of GQDs. (**b**) The measured OD600 of bacteria over time in the presence of GQDs (10 _μ_g/mL), hydrogen peroxide (HP, 4.5 mM), or both simultaneously. The data is presented as either a bar chart (left) or a scatter plot (right). The bars on the left and data points on the right represent average values. In the bar charts, data points represent individual measurements acquired from replicate experiments. All error bars represent the standard deviation of the mean.

We next examined whether GQDs alter bacterial growth in the presence of exogenous H_2_O_2_ (Figure. 1b). As expected, H_2_O_2_ alone suppressed early culture growth by extending the lag phase. In contrast, cultures treated with both H_2_O_2_ and GQDs showed improved growth over the first 6 hours, indicating that GQDs mitigate the early growth inhibition caused by peroxide exposure. After overnight growth, the GQD + H_2_O_2_ condition reached a final OD600 comparable to the GQD-only condition, whereas the H_2_O_2_-only condition remained lower. Thus, although peroxide delays the onset of growth, GQDs substantially blunt this effect and promote recovery to near-control densities in the stationary phase. Similar behavior was also observed at higher GQD concentrations of 25 and 50 _μ_g/mL (Figure. S1). Collectively, these result show that GQDs do not impair *E. coli* growth under basal conditions and help preserve bacterial growth under peroxide-induced oxidative stress.

### 3.2. GQDs attenuate oxidation in bacterial membranes

To determine whether GQDs influence oxidative damage in bacterial membranes, we used C11-BODIPY, a membrane-partitioning fluorescent probe that undergoes a blue-shift in its emission maximum from 590 nm to 520 nm upon oxidation.^57^ Figure 2a shows the structure of C11-BODIPY and a schematic of its partitioning into bacterial membranes following incubation with *E. coli*. Representative confocal fluorescence images confirmed successful incorporation of the probe into bacteria, as fluorescence was clearly detected in both the 590 nm and 520 nm channels (Figure. 2b). These results establish that C11-BODIPY was effectively associated with the bacterial membrane environment and could be used to monitor oxidation under the tested conditions.

**Figure 2.**
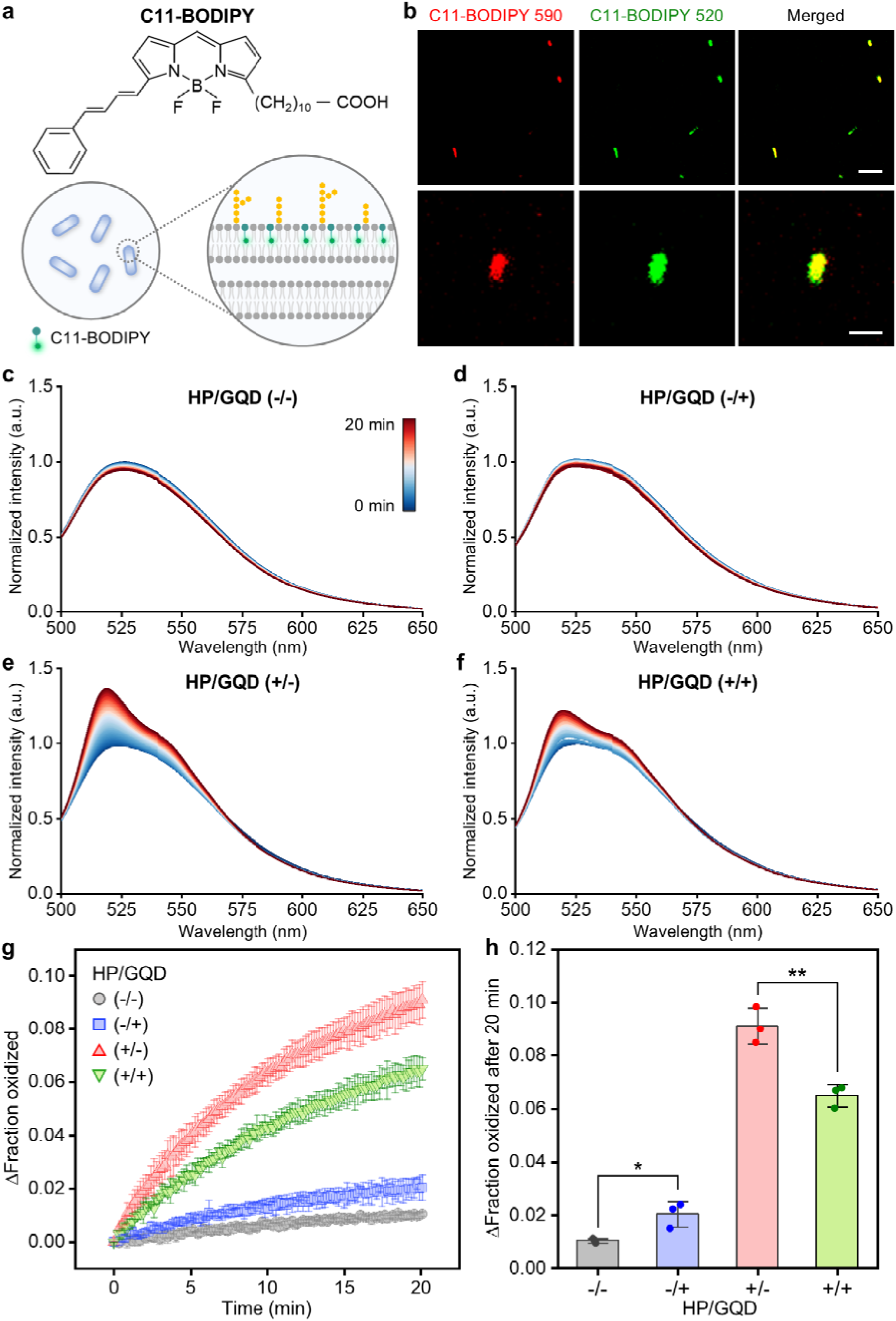
GQDs inhibit oxidation of C11-BODIPY within the biological membranes of *E. coli*. (**a**) Structure of C11-BODIPY and schematic of its partitioning into bacterial membranes. (**b**) Representative confocal fluorescence images of bacteria that were labeled with C11-BODIPY. Scale bars represent 10 μm (top) and 2 μm (bottom). Representative time-dependent fluorescence emission spectra of C11-BODIPY-labeled bacteria in (**c**) untreated conditions, (**d**) the presence of GQDs, (**e**) the presence of HP, or (**f**) the presence of GQDs and HP simultaneously. The fractional increase in oxidized C11-BODIPY (**g**) over time and (**h**) after 20 minutes in either the presence or absence of HP or GQDs. All graphs in (c)-(f) are normalized to the maximum intensity of the initial spectrum. All HP and GQD concentrations in (d)-(h) are 4.5 mM and 50 μg/mL, respectively. In (g), data points represent the average of triplicate measurements and error bars represent the standard deviation. In (h), bars represent the average value, error bars represent the standard deviation, and data points represent the individual replicate measurements (N=3; unpaired, two-sample, t-test; * = P<0.05, ** = P<0.01).

We next used fluorescence spectroscopy to monitor the time-dependent emission spectra of C11-BODIPY under each condition. In untreated samples, the spectra changed minimally over the 20-minute observation window (Figure. 2c). Similarly, only small spectral changes were observed in the presence of GQDs alone (Figure. 2d), indicating that GQDs have little effect on C11-BODIPY oxidation in the absence of added peroxide. In contrast, addition of H_2_O_2_ produced a pronounced increase in emission near 520 nm, consistent with oxidation of the probe (Figure. 2e). A similar shift was observed when bacteria were exposed to both H_2_O_2_ and GQDs (Figure. 2f), but the change at 520 nm was clearly less pronounced, suggesting that GQDs attenuate peroxide-induced membrane oxidation.

Importantly, control experiments performed using bacterial media with and without H_2_O_2_ showed no measurable spectral changes over the same 20-minute interval (Figure. S2), indicating that the changes observed in Figure 2 do not arise from media autofluorescence. To quantify these spectral changes, we converted the emission spectra in Figures 2c–2f into oxidized C11-BODIPY fractions using the calibration procedures described in the Supporting Information. This analysis yielded the change in the oxidized fraction of C11-BODIPY as a function of time, shown in Figure 2g. Consistent with the qualitative spectral trends, untreated bacteria exhibited only a small increase over time, while bacteria exposed to GQDs alone showed a slightly larger increase. This minor increase could be attributed to the potential generation of ROS by GQDs upon constant irradiation by excitation visible light throughout the fluorescence measurement.^42, 58^ The response induced by H_2_O_2_ was significantly greater, which produced a rapid rise in the change of oxidized fraction approaching 10% over 20 minutes. In the presence of both H_2_O_2_ and GQDs, the increase remained substantial but was markedly reduced relative to conditions in which bacteria were exposed to H_2_O_2_ alone.

This protective effect is also evident from the endpoint comparison after 20 minutes (Figure. 2h). Relative to untreated bacteria, H_2_O_2_ caused a large increase in C11-BODIPY oxidation, whereas inclusion of GQDs significantly reduced this peroxide-driven response. Thus, while GQDs alone produce little change in membrane oxidation over this time frame, they substantially suppress the much larger oxidative damage induced by exogenous peroxide. Together, these results indicate that GQDs attenuate H_2_O_2_-induced oxidation in bacterial membranes.

### 3.3. GQDs suppress intracellular ROS accumulation and oxidative stress in bacteria

To probe intracellular oxidative stress in bacteria, we transformed *E. coli* to express the H_2_O_2_-responsive redox sensor roGFP2-Orp1 (Figure. 3a).^50^ In this construct, Orp1 reacts directly with H_2_O_2_ and transfers this oxidation to roGFP2, which alters the excitation properties of the fluorophore.^50, 59, 60^ Oxidation of the sensor is associated with an increase in the relative fluorescence excitation signal at 405 nm and a decrease at 488 nm.^61^ Thus, an increase in the 405 nm/488 nm ratio reports greater intracellular H_2_O_2_ accumulation.

**Figure 3.**
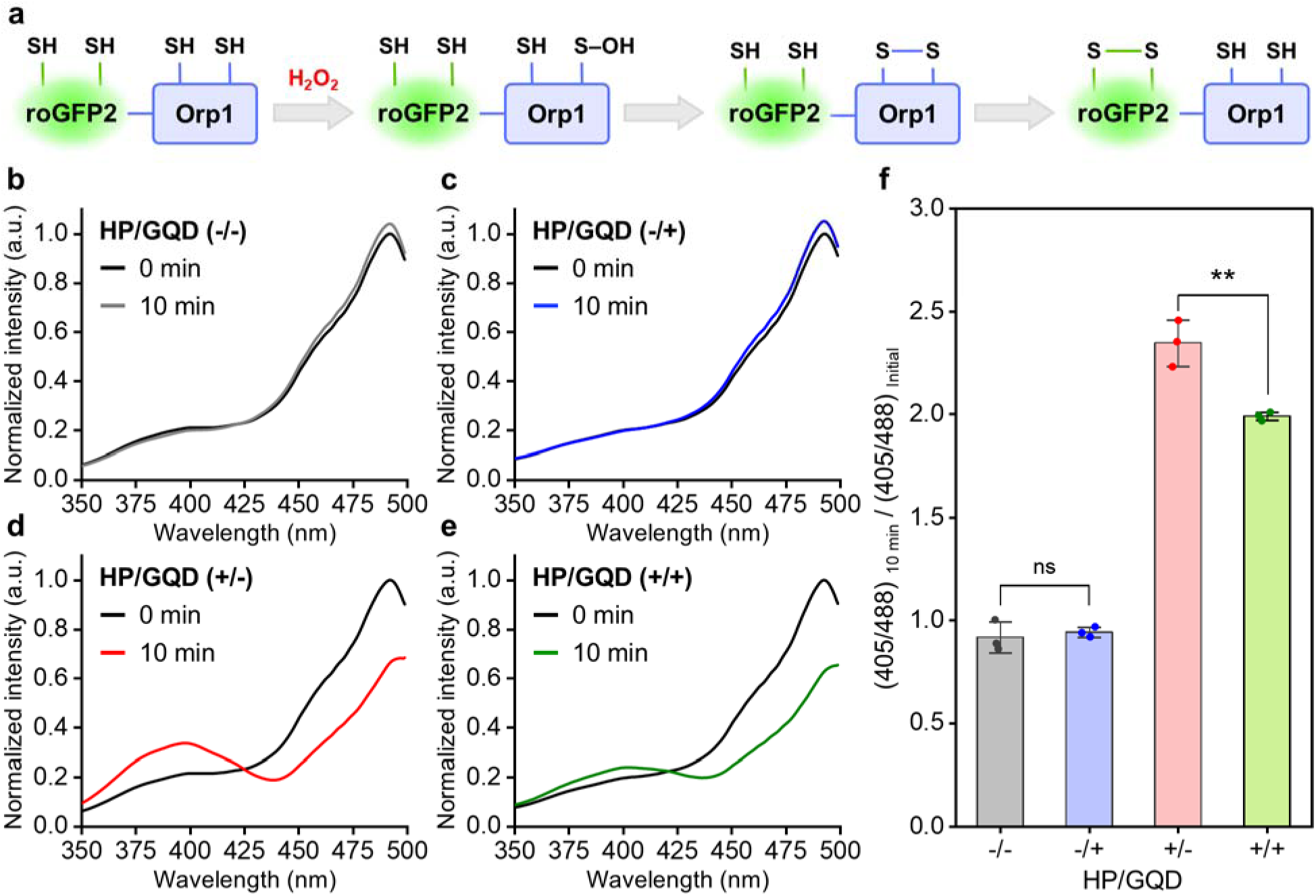
GQDs inhibit intracellular accumulation of H_2_O_2_ in *E. coli*. (**a**) Schematic of the H_2_O_2_ reaction sequence with roGFP2-Orp1. Normalized excitation spectra of bacteria expressing roGFP2-Orp1 in (**b**) untreated conditions, (**c**) the presence of GQDs, (**d**) the presence of hydrogen peroxide (HP), (**e**) or the presence of GQDs and HP simultaneously. (**f**) The relative change in the 405 nm/488 nm roGFP2-Orp1 excitation ratio after 10 minutes, either in the presence or absence of HP or GQDs. All HP and GQD concentrations in (c)-(f) are 0.1 mM and 50 _μ_g/mL, respectively. In (f), bars represent the average value, error bars represent the standard deviation, and data points represent the individual replicat measurements (N=3; unpaired, two-sample, t-test; ns = non-significant, ** = P<0.01).

In untreated bacteria, the normalized excitation spectra changed only minimally over the 10-minute measurement window (Figure. 3b). Similarly, little change was observed when bacteria were exposed to GQDs alone (Figure. 3c), as expected since roGFP2-Orp1 responds directly to H_2_O_2_. Therefore, the absence of a strong spectral shift indicates that GQDs do not themselves promote appreciable intracellular peroxide accumulation in this experimental condition. Importantly, control measurements with GQDs in the absence of bacteria confirmed that changes in GQD excitation were minimal over this 10-minute time interval (Figure. S3), thus the results were not obscured by GQD spectral overlap. Incubation with H_2_O_2_ produced a clear shift in the normalized excitation spectrum, characterized by an increase in the relative 405 nm signal and a corresponding decrease in the relative 488 nm signal (Figure. 3d). This shift indicates substantial oxidation of roGFP2-Orp1 due to accumulation of peroxide within the bacteria. When GQDs were present together with H_2_O_2_, the same overall trend was observed, but the spectral shift was clearly less pronounced (Figure. 3e).

This behavior is captured in the endpoint quantification of the 405 nm/488 nm ratio after 10 minutes (Figure. 3f). Relative to untreated bacteria, H_2_O_2_ caused a strong increase in the roGFP2-Orp1 oxidation response, whereas the presence of GQDs significantly attenuated this increase. These results indicate that GQDs suppress intracellular H_2_O_2_ accumulation in *E. coli*. A likely explanation is that GQDs interact with and sequester ROS in the bulk solution before they can enter the cell, thereby lowering the oxidative burden experienced intracellularly. This interpretation is consistent with previously reported ROS scavenging behavior of GQDs in biological systems.^35, 36, 39^

We next asked whether the reduction in intracellular H_2_O_2_ accumulation also translated to reduced disruption of intracellular redox homeostasis. To probe this, we transformed *E. coli* to express Grx1-roGFP2, a glutathione-responsive sensor that reports on the redox state of the glutathione/glutaredoxin system (Figure 4a).^51, 60, 62^ Glutathione is an endogenous antioxidant that neutralizes excess ROS by forming glutathione disulfide bonds on proteins to prevent irreversible protein damage.^63, 64^ The glutathionylated disulfide substrate can then be reduced back through glutaredoxin.^65^ In Grx1-roGFP2, oxidation within the glutathione pool is relayed through Grx1 to roGFP2, producing an increase in the 405 nm/488 nm ratio in the excitation spectrum. Thus, whereas roGFP2-Orp1 reports on intracellular H_2_O_2_ accumulation, Grx1-roGFP2 reports on the downstream oxidative perturbation experienced by a major intracellular redox buffering system.

**Figure 4.**
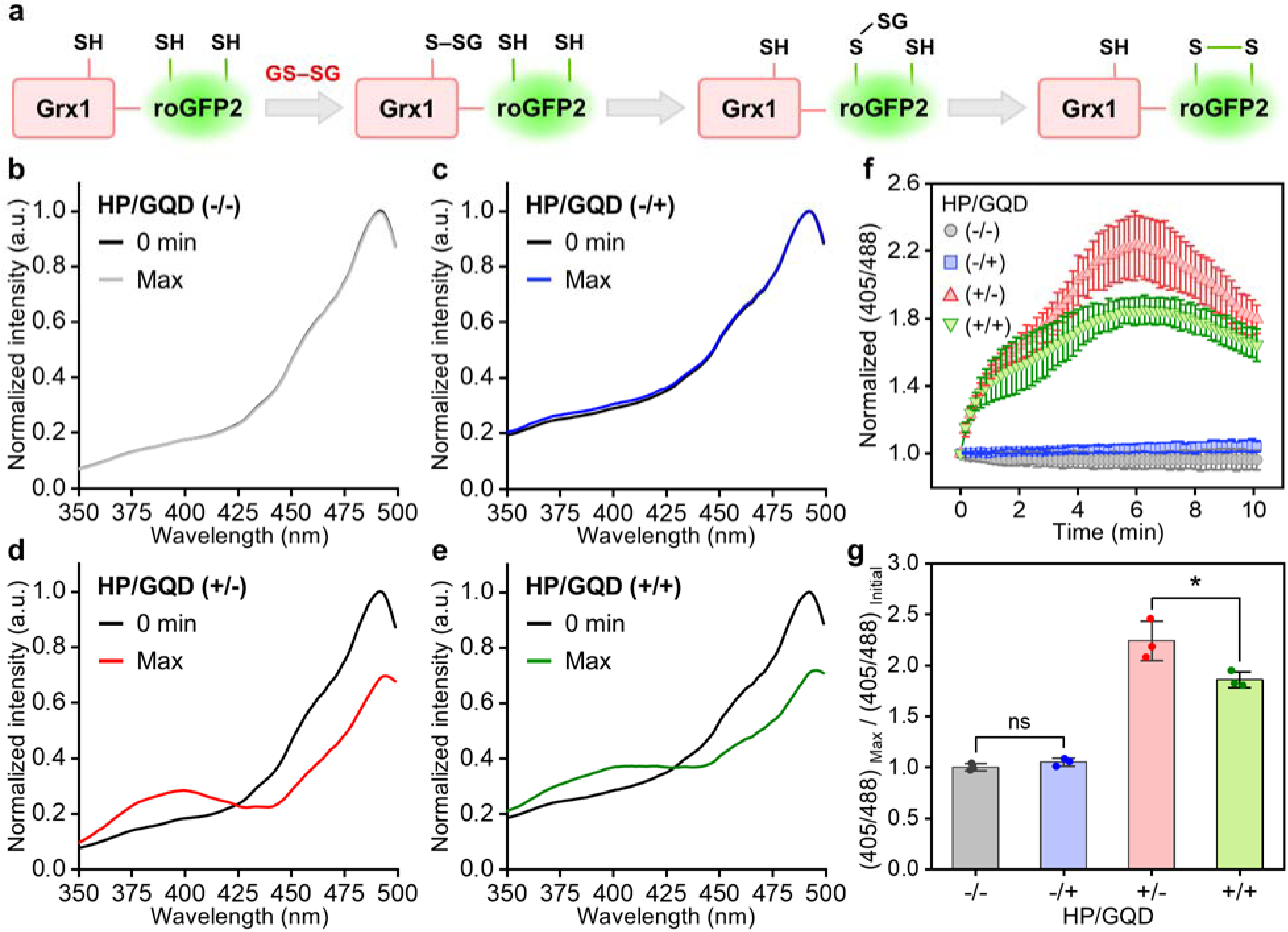
GQDs attenuate oxidative stress-induced disruption of glutathione redox homeostasis in *E. coli*. (**a**) Schematic of the glutathione disulfide (GS-SG) reaction sequence with Grx1-roGFP2. Normalized excitation spectra of bacteria expressing Grx1-roGFP2 in (**b**) untreated condition, (**c**) the presence of GQDs, (**d**) the presence of hydrogen peroxide (HP), or (**e**) the presence of GQDs and HP simultaneously. In (b)-(e), “Max” refers to the maximum spectral shift observed over a 10-minute interval. The normalized 405 nm/488 nm Grx1-roGFP2 excitation ratio (**f**) over 10 minutes and (**g**) at its maximal value. All HP and GQD concentrations in (c)-(g) are 0.5 mM and 50 _μ_g/mL, respectively. In (f), data points represent the average of triplicate measurements and error bars represent the standard deviation. In (g), bars represent the average value, error bars represent the standard deviation, and data points represent the individual replicate measurements (N=3; unpaired, two-sample, t-test; ns = non-significant, * = P<0.05).

Consistent with the roGFP2-Orp1 measurements, untreated bacteria exhibited minimal spectral change over the measurement window (Figure. 4b), and the presence of GQDs alone also produced little change (Figure. 4c). In contrast, exposure to H_2_O_2_ caused a pronounced shift in the normalized excitation spectrum, characterized by an increase in the relative 405 nm signal and a corresponding decrease in the relative 488 nm signal (Figure. 4d), indicating substantial oxidation within the glutathione redox network. When bacteria were exposed to both H_2_O_2_ and GQDs, the same general shift was still observed, but its magnitude was reduced (Figure. 4e).

Notably, the spectra in Figures 4b–4e are shown at the point of maximal spectral change observed during the 10-minute interval. This is because the response in this system exhibited a pronounced spike that reached a local maximum after approximately 5 to 6 minutes (Figure. 4f), rather than a sustained value over time as observed with roGFP2-Orp1. This profile suggests that peroxide exposure produces a rapid but partially recoverable perturbation to glutathione redox homeostasis. In the presence of H_2_O_2_ alone, this spike was large, whereas in the presence of both H_2_O_2_ and GQDs, the maximum response was clearly reduced. Because the largest difference between conditions occurred at the peak of this transient response, we quantified the maximal 405 nm/488 nm ratio rather than the endpoint value (Figure. 4g). This analysis showed that H_2_O_2_ caused a strong increase in the maximal oxidation response of Grx1-roGFP2, while inclusion of GQDs significantly attenuated this increase.

Together, these results show that GQDs not only reduce intracellular H_2_O_2_ accumulation, but also lessen the downstream oxidative disruption imposed on the glutathione/glutaredoxin system. This behavior is consistent with the idea that GQDs suppress peroxide-driven oxidative stress before it can propagate into broader intracellular redox imbalance.

## 4. CONCLUSION

In conclusion, we show that GQDs containing carboxyl and hydroxyl functional groups mitigate peroxide-induced oxidative stress in *E. coli* without adversely affecting bacterial growth under basal conditions. Using a myriad of complementary spectroscopic techniques, we found that these GQDs preserve bacterial growth in the presence of exogenous H_2_O_2_, attenuate oxidation in bacterial membranes, suppress intracellular H_2_O_2_ accumulation, and lessen oxidative disruption of intracellular redox buffering networks. Taken together, these results support a model in which these types of GQDs act upstream of intracellular oxidative damage by limiting peroxide-driven oxidative stress before it propagates across the bacterial envelope and into broader redox imbalance. More broadly, this work expands understanding of how graphene-based nanomaterials interact with microbial systems under oxidative stress and points to carboxyl/hydroxyl-functionalized GQDs as promising additives for formulations in which protection of microbial cargo from oxidative damage is important. These findings warrant further investigation of such roles in microbial preservation, delivery, and related biotechnological applications.

## ASSOCIATED CONTENT

## Supporting Information

The Supporting Information is available free of charge at [link TBD]

- Calculation of the molar fraction of oxidized C11-BODIPY; Figure S1. GQDs inhibit the suppressed growth rate of *E. coli* under oxidative stress; Figure S2. Media autofluorescence does not change significantly over time; Figure S3. Excitation intensity of GQDs does not change significantly after 10 minutes of incubation (PDF).

## Author Contributions

W.F.Z. and J.K. designed the work. J.K. and S.N.B conducted experiments. All authors contributed to analyzing data and preparing the manuscript.

## Notes

The authors declare no competing interests.

## ACKNOWLEDGEMENTS

This work was supported in part by the National Institutes of Health through R35GM147333 to J. Kim and W. F. Zeno.

## Notes

### Competing Interest Statement

The authors have declared no competing interest.

