## Supporting Information for "Graphene Quantum Dots Mitigate Oxidative Stress in Bacteria"

**Calculation of the molar fraction of oxidized C11-BODIPY**

Emission intensity of oxidized and unoxidized state of C11-BODIPY follows equations 1 and 2, respectively. The ratio of fluorescence intensity coefficients, R (eq 3), was determined from average coefficient ratios calculated from membranes of various compositions,[1] as microbial membranes are composed of various types of phospholipids.[2] Finally, the molar fraction of oxidized C11-BODIPY was calculated using equation 4.

E: fluorescence intensity at 520 nm and 590 nm

ε: fluorescence intensity coefficient of 520 nm and 590 nm

C: concentration of oxidized and unoxidized C11-BODIPY

$$\mathrm{E520} = \varepsilon 520 \times \mathrm{C_{ox}} \qquad \text{eq (1)}$$

$$\mathrm{E590} = \varepsilon 590 \times \mathrm{C_{unox}} \qquad \text{eq (2)}$$

$$\mathrm{R} = \frac{\varepsilon 520}{\varepsilon 590} \qquad \text{eq (3)}$$

$$\text{Fraction oxidized} = \frac{\mathrm{C_{ox}}}{\mathrm{C_{ox}} + \mathrm{C_{unox}}} = \frac{\mathrm{E520}/\varepsilon 520}{\mathrm{E520}/\varepsilon 520 + \mathrm{E590}/\varepsilon 590} = \frac{\mathrm{E520}}{\mathrm{E520} + \mathrm{R} \times \mathrm{E590}} \qquad \text{eq (4)}$$

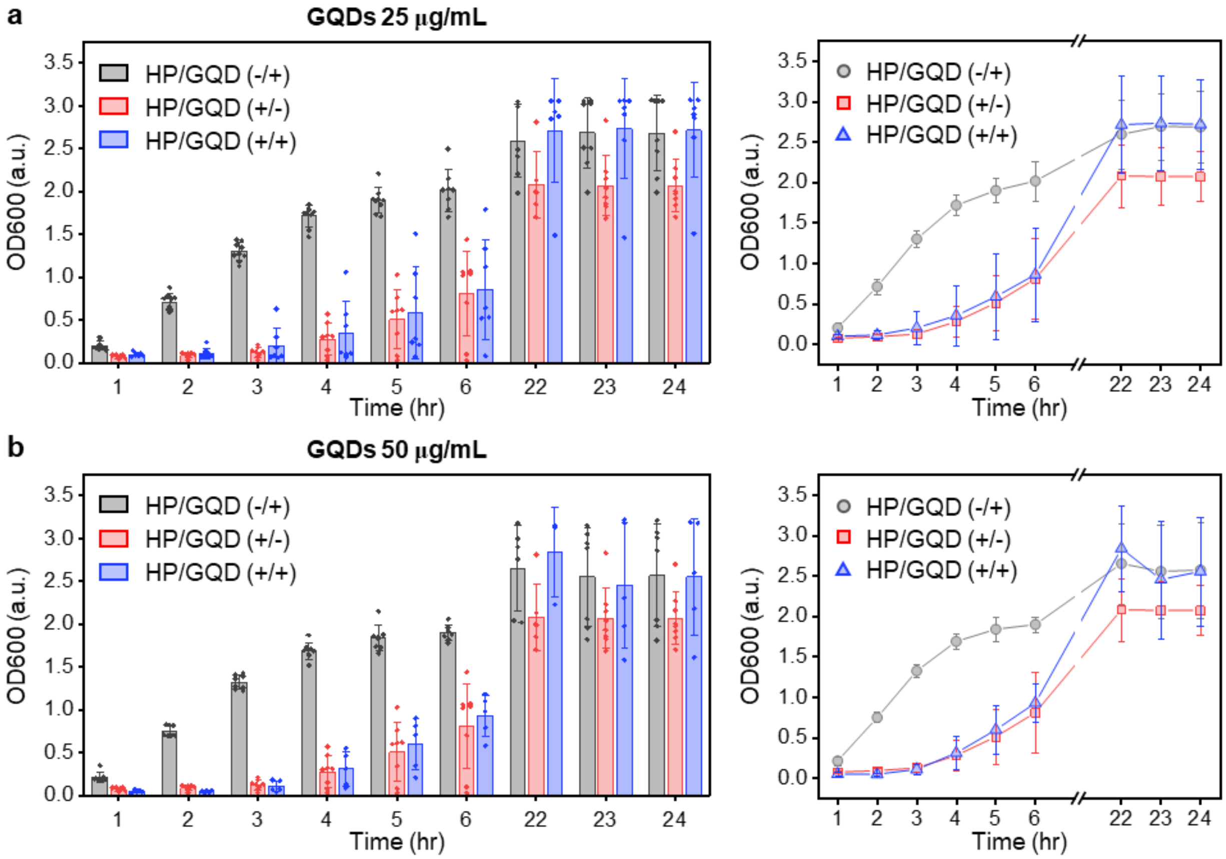


**Figure S1. GQDs inhibit the suppressed growth rate of *E. coli* under oxidative stress.** The measured OD600 of bacteria over time in the presence of GQDs, hydrogen peroxide (HP, 4.5 mM), or both simultaneously. The final concentration of GQDs was (**a**) 25 μg/mL and (**b**) 50 μg/mL. The data is presented as either a bar chart (left) or a scatter plot (right). The bars on the left and data points on the right represent the average values. In the bar charts, data points represent individual measurements acquired from replicate experiments. All error bars represent the standard deviation of the mean.

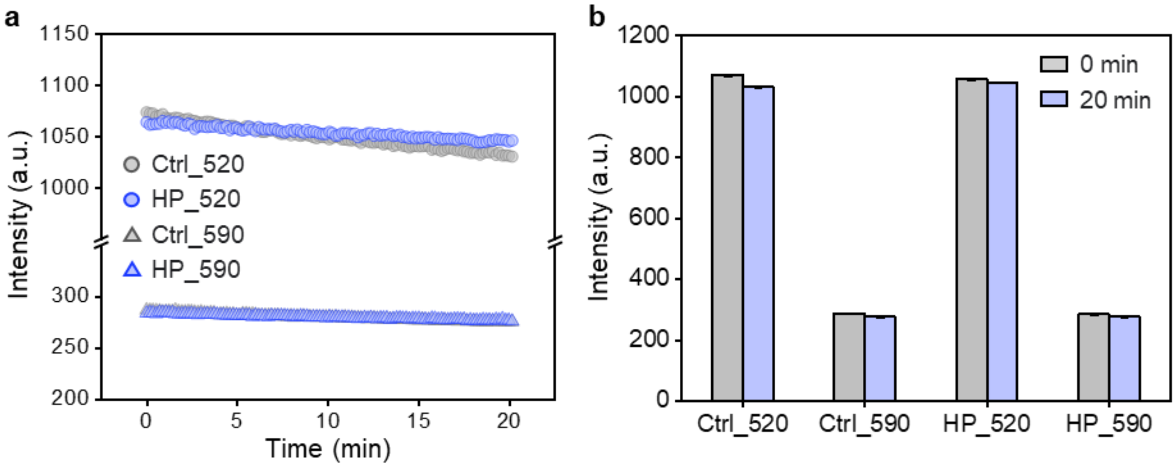


**Figure S2. Media autofluorescence does not change significantly over time.** (**a**) The emission intensity of media at 520 nm and 590 nm with and without $H_2O_2$. Ctrl refers to the media without $H_2O_2$, and HP refers to the media with 4.5 mM $H_2O_2$. (**b**) The average emission intensity of initial measurements (0 min) and after 20 minutes of incubation with and without $H_2O_2$. The emission intensity was acquired every 10 seconds for 20 minutes at 37 °C. Bars in (b) represent the average emission intensity of the initial and last three measurements, and error bars represent the standard deviation.

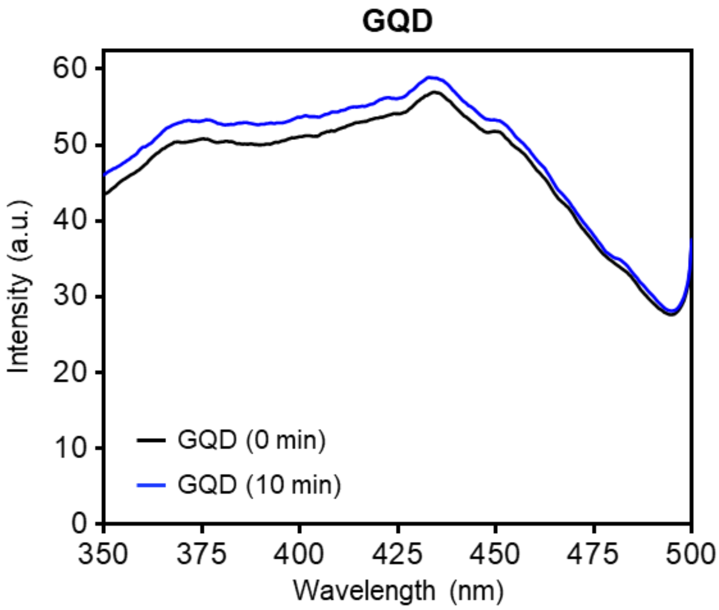


**Figure S3. Excitation intensity of GQDs does not change significantly after 10 minutes of incubation.** Representative excitation spectra of 50 μg/mL GQD before and after incubation at 25 $^{o}$C for 10 minutes at the emission wavelength of 510 nm.
